# The attachment of Sso7d-like protein improves processivity and resistance to inhibitors of M-MuLV reverse transcriptase

**DOI:** 10.1101/2020.07.02.185637

**Authors:** Igor P. Oscorbin, Pei Fong Wong, Ulyana A. Boyarskikh, Evgeny A. Khrapov, Maksim L. Filipenko

**Affiliations:** Institute of Chemical Biology and Fundamental Medicine, Siberian Branch of the Russian Academy of Sciences, Lavrentiev av. 8, Novosibirsk, 630090, Russian Federation; Novosibirsk State University, Pirogova str. 2, Novosibirsk, 630090, Russian Federation

**Keywords:** reverse transcription, cDNA synthesis, M-MuLV reverse transcriptase, chimeric protein, protein engineering

## Abstract

Reverse transcriptases, RTs, are a standard tool in both fundamental studies and diagnostics used for transcriptome profiling, virus RNA testing and other tasks. RTs should possess elevated temperature optimum, high thermal stability, processivity, and tolerance to contaminants originating from the biological substances under analysis or the purification reagents. Here, we have constructed a set of chimeric RTs, based on the combination of MuLV-RT and DNA-binding domains: the DNA-binding domain of DNA ligase *Pyrococcus abyssi* and Sto7d protein, Sso7d counterpart, from *Sulfolobus tokodaii*. Chimeric RTs showed the same optimal temperature and the efficacy of terminal transferase reaction as the original M-MuLV RT. Processivity and the efficiency in cDNA synthesis of the chimeric RT with Sto7d at C-end were increased several-fold. The attachment of Sto7d enhanced the M-MuLV RT tolerance to the most common amplification inhibitors: NaCl, urea, guanidinium chloride, formamide, components of human whole blood, and human blood plasma. Thus, fusing M-MuLV RT with an additional domain resulted in more robust and efficient RTs.

## INTRODUCTION

Reverse transcriptases (RTs), discovered in 1970, belong to a subfamily of DNA polymerases and are capable of synthesising DNA using RNA as a template (Coffin and Fan, 2016). After decades of studies, a significant number of RTs has been discovered in all three domains of life, and several of them have been implemented into practical applications (Efstratiadis et al., 1975; Kotewicz et al., 1988). *In vivo*, RTs participate in a lifecycle of retroviruses, take part in retrotransposon expansion, and serve as telomerase elongating linear chromosome ends in eukaryotes. Given their unique ability to produce complementary DNA, cDNA, RTs became one of the crucial tools in both fundamental studies and diagnostics, where RNA is a matter of concern.

The most commonly used enzymes are reverse transcriptases from Moloney murine leukaemia virus and avian myeloblastosis virus (M-MuLV and AMV RTs, respectively). However, biochemical characteristics of M-MuLV and AMV RTs are far from ideal. Both enzymes have a thermal optimum at 37-45□C, display relatively low processivity and tolerance to inhibitors, and have no proofreading activity (Gerard et al., 2002; “Polymerization and RNase H activities of the reverse transcriptases from avian myeloblastosis, human immunodeficiency, and Moloney murine leukemia… - PubMed - NCBI,” n.d.; Yasukawa et al., 2008). The conversion of complicated structured RNA templates to cDNA requires the elevated reaction temperature about 6□C to denature the template obstacles that hinder RTs catalysis (Lanciault and Champoux, 2006). Traces of contaminants originating from test specimens or purification reagents can lead to false-negative results when testing infections in complex biological substances, like blood, sputum, faeces. In addition, the low fidelity of cDNA synthesis makes it challenging to analyse somatic mutations, especially in initially degraded RNAs, purified from formalin-fixed paraffin-embedded (FFPE) specimens that are an invaluable source of information when treating oncological diseases.

Multiple attempts have been made to obtain novel more processive, accurate, and resistant to inhibitors RTs. Two main strategies have been used to achieve this goal: random or site-directed mutagenesis of already known RTs and characterisation of novel RTs. The first approach was successfully used for improving M-MuLV and AMV RTs, resulting in more thermal stable enzymes (Arezi and Hogrefe, 2009; Baba et al., 2017; Baranauskas et al., 2012a; Katano et al., 2017; Konishi et al., 2012; Yasukawa et al., 2010). While being weak and requiring manganese ions (Griffiths et al., 2004; Myers and Gelfand, 1991), innate reverse transcriptase activity of family A DNA polymerases can be enhanced using unusual mutagenesis (Sano et al., 2012; Sauter and Marx, 2006) or fusing with protein domains from other DNA polymerases to enhance the enzyme processivity (Schönbrunner et al., 2006). Moreover, thermostable family B DNA polymerases possessing both proofreading and RT activity were created through random mutagenesis of KOD DNA-polymerase from *Thermococcus gorgonarius* and *Thermococcus kodakaraensis* KOD1, respectively (Ellefson et al., 2016; Jozwiakowski and Connolly, 2011). By using the second strategy, the novel RTs with enhanced thermal stability and processivity have been recruited from metagenomes (Heller et al., 2019; Michael J Moser et al., 2012) and bacterial type II introns (Blocker et al., 2005; Mohr et al., 2013; Ng et al., 2007; Saldanha et al., 1999; Vellore et al., 2004; Zhao et al., 2018a).

Despite the progress achieved in the field, previous endeavours have mainly focused on obtaining enhanced thermostability, leaving other the other enzymatic properties in the shadow. Thus, only a few works have described increasing processivity (Baranauskas et al., 2012b), tolerance to inhibitors (Arezi et al., 2010a) or emerging of proofreading activity (Ellefson et al., 2016). In terms of practical usage, especially for diagnostic purposes, the tolerance to inhibitors is necessary for the development of reliable tests. Therefore, the need for an enzyme with improved applicability is still of high relevance.

Fusion with an additional protein domain is a convenient and promising technique for constructing enzymes with improved characteristics. Previously, this approach has been successfully applied to improve Pfu-, Taq-, and Φ29 polymerases (de Vega et al., 2010; Oscorbin et al., 2017; Pavlov et al., 2012, 2002; Wang et al., 2004). Specifically, the DNA-binding protein Sso7d from thermophilic archaea *Sulfolobus solfataricus* was attached to Pfu-polymerase; the DNA-binding domain of TopoV of *Methanopyrus kandleri* was attached to Φ29 polymerase. We have characterised an improved Bst-like Gss-polymerase with Sto7d protein in C-terminus; the enzyme showed enhanced processivity and tolerance to amplification inhibitors. However, to our knowledge, no chimeric RTs with additional domains were designed. In the present work, we constructed a set of chimeric reverse transcriptases based on MuLV-RT and DNA-binding domains: the DNA-binding domain of DNA ligase *Pyrococcus abyssi* and Sto7d protein, a counterpart of Sso7d from *Sulfolobus tokodaii*.

## MATERIALS AND METHODS

### Construction of M-MLV reverse transcriptase fusions

The set of fusion proteins was constructed: M-MuLV-RT with the DBD of ATP-dependent DNA ligase from *Pyrococcus abyssi* at N- or C-terminus, M-MuLV-RT with Sto7d protein from *Sulfolobus tokodaii* at N- or C-terminus, M-MuLV with D200N, T330P, L139P mutations, previously characterised as enhancing the affinity to an RNA template (Arezi and Hogrefe, 2009), and M-MuLV with D200N, T330P, L139P with Sto7d protein at C-terminus. Coding sequences for all enzymes were cloned into pET23b (Novagen, USA) using overlapping PCR and REase-ligation methodology, as fully described in the Supplementary file.

### Expression and purification of reverse transcriptases

BL21 (DE3) pLysS (Promega, USA) strain of *E. coli* cells harbouring the plasmid, encoding RT, was grown to OD_600_ = 0.3 in LB medium at 37 °C. Four litres of LB in a LiFlus GX fermenter (Biotron Inc., South Korea) were inoculated with 40 ml of the previously obtained culture, and the cells were grown to OD_600_ = 0.6 at 37 °C. Expression was induced by the addition of IPTG up to 1 mM concentration. After induction for 4 h at 37 °C, the cells were harvested by centrifugation at 4,000 × *g* and stored at −70 °C. The cell pellets were resuspended in the lysis buffer (50 mM T ris-HCl pH 7.0, 2.5 mM MgCl_2_, 0.1 mM CaCl_2_, 1 mM PMSF, 1 mg/mL lysozyme), incubated for a 30 min on ice followed by sonication. The lysates were treated for 15 min at 37 °C with DNAse I (1 μg/ml) for DNA digestion followed by centrifugation at 14,000 × *g*. The resulting supernatants were loaded onto a 5-ml IMAC (Bio-Rad, USA) column pre-equilibrated with buffer A (50 mM Tris-HCl pH 7.0, 0.3 M NaCl), followed by washing the column with 25 ml of buffer A. Bound proteins were eluted using buffer B (Buffer A with 0.3 mM imidazole). After affinity chromatography, the fractions that contained RTs were pooled and loaded onto a 2-ml Macro-Prep DEAE Resin (Bio-Rad, USA) column pre-equilibrated with buffer C (50 mM Tris-HCl, 0.1 mM EDTA, pH 7.5). The column was washed with 10 ml of buffer C, and bound proteins were eluted by a 0-100% linear gradient of buffer D (50 mM Tris-HCl, 1 M NaCl, 0.1 mM EDTA, pH 7.5). Fractions with RTs were pooled, dialyzed against the storage buffer (50 mM Tris-HCl, 150 mM NaCl, 0.1 mM EDTA, 50% Glycerol, 0.1% NP-40, pH 7.5), and stored at −20 °C. All fractions from each step were analysed by SDS-PAGE. The purity of the preparations was not less than 95%. The concentration of purified proteins was measured using a standard Bradford assay.

### RNA-dependent DNA-polymerase activity measurement

The specific activity of RT was assayed by using radiolabelled nucleotides incorporation. The reaction mix (50 μL) contained 0.4 mM poly(rA)/oligo(dT)_25_ (concentration defined by oligo(dT)_25_), 0.5 mM α-[^32^P]-dTTP (4 Bq/pmol), 50 mM Tris-HCI (pH 8.3), 6 mM MgCl_2_, 10 mM dithiothreitol. The reactions were initiated by adding the enzyme on ice; the samples were immediately transferred to a preheated thermal cycler for incubation at 37 °C for 10 min followed by inactivation by heating at 90°C for a 1 min. The reaction products were collected on DE81 paper (Sigma-Aldrich Corp.), washed twice with 0.5 M Na_2_HPO_4_, and counted in a Pharos PX (Bio-Rad, USA). One unit of polymerase activity was defined as the amount of enzyme that incorporated 1 nmol of dTTP into acid-insoluble material in 10 minutes at 37°C.

### Thermal stability, optimal temperature and ion concentration

The thermal stability of enzymes was studied by heating the enzymes followed by the polymerase activity assay. The aliquots of polymerase activity reaction buffer (described above) containing an identical amount (0.2 U) of the enzymes were incubated at 40, or 50°C for 15-60, or 5-15 min, respectively. Optionally, 0.8 mM the primed substrate (poly(rA)/oligo(dT)_25_) was added in the heating reaction. The reactions were chilled on ice, and the polymerase activity was measured as mentioned above.

The temperature optimum of the enzymes was investigated by measuring polymerase activity at a range of 30-60°C by 5°C per step, while other conditions were identical to those described above for the polymerase activity assay.

The optimal ion concentrations were examined similarly as in the polymerase activity assay using concentrations of K_2_SO_4_, NH_4_Cl, KCl, (NH_4_)_2_SO_4_ in the range of 50–500 mM.

### Thermal shift assay

The thermal stability of the enzymes was analysed by heating in a CFX96 Touch™ Real-Time PCR Detection System (Bio-Rad, USA). The reactions were conducted in 20 μL containing 1× enzyme storage buffer (50 mM Tris-HCl, 150 mM NaCl, 0.1 mM EDTA, 50% glycerol, 0.1% NP-40, pH 7.5), 1× SYPRO Orange (Invitrogen, USA) and 2 μM of enzymes. Optionally, 0.4 mM poly(rA)•oligo(dT)_25_ (concentration defined by oligo(dT)_25_) was added to the reactions. The temperature was increased from 25 to 70°C, with an increment of 0.3°C per 20 s and collection of the fluorescent signal in FRET mode.

### RNA purification from FFPE samples

FFPE samples for RNA purification were collected from 10 colon cancer patients who had been operated in Novosibirsk regional cancer centre. Informed consent was obtained from all individual participants involved in the study. All the research procedures involving human participants were performed in accordance with the ethical standards of the institutional and national research committee and the 1964 Helsinki declaration and its later amendments or comparable ethical standards. FFPE sections (10 μm) were deparaffinised by heating for 3 min at 80°C with 300 μL of mineral oil (ICN, USA). 200 μL of Proteinase K solution (1% SDS, 10 mM EDTA, 100 mM NaCl, 10 mM Tris-HCl pH8.0, 1 mg/mL Proteinase K (SibEnzyme, Russia)) was added to each sample, followed by overnight digestion at 56°C. The RNA was isolated from digested specimens using a standard phenol-chloroform extraction followed by isopropanol precipitation. The residual DNA traces were removed by the DNAse I treatment (Worthington Corporation, USA) followed by the second round of phenol-chloroform extraction and isopropanol precipitation. The purified RNA was dissolved in DEPC-treated water and stored at −80°C.

### In vitro RNA synthesis

Linearised plasmid pBlueScript with 3 kbp fragment of lambda phage genome (positions 6499-9543, GenBank ID J02459.1) inserted at EcoRI/HindIII sites was used as the template for RNA synthesis. The reactions were conducted in a 50 μL volume containing 2 μg of DNA template, digested by EcoRI, 1 mM NTP, 100 U of T7 RNA polymerase (SibEnzyme, Russia), 1× reaction buffer (50 mM Tris-HCl, pH 7.5, 6 mM MgCl_2_, 10 mM DTT, 2 mM spermidine). After mixing, the probes were incubated for 2 hours at 37°C, followed by DNAse I treatment (100 U, 15 minutes at 37°C) and RNA purification with phenol-chloroform extraction and isopropanol precipitation. The purified RNA was dissolved in DEPC-treated water and stored at −80°C.

### MS2 phage RNA purification

The MS2 phage was grown using a modified protocol from Sambrook and Russel (Evans, 1990). The overnight culture of *E. coli* strain K12 was diluted in 3 mL of MS2 broth to a final OD_600_ = 1 (1 ×10^9^ cells/mL), followed by the addition of MS2 phage to achieve a multiplicity of infection of 5. The culture was incubated at 37°C for 20 min, mixed with 500 mL of prewarmed MS2 broth, and incubated for 12 hours. Cell lysis was induced by the addition of 20 ml of chloroform, followed by shaking for 10 min at 37°C. The lysate was treated with DNAse I and RNAse A (50 mg/mL) for a 30 min at 37°C, then NaCl was added to a final concentration of 1 M. The mixture was incubated on ice for a 1 hour, followed by centrifugation (10 min, 11000×*g*) at 4°C. The ammonium sulfate was added to the supernatant (to 20% final concentration (w/w)), and the mixture was incubated for 2 hours at 4°C, followed by centrifugation (30 min, 11000×*g*) at 4°C. The extra ammonium sulfate was added to the supernatant (to 50% final concentration (w/w)), and the mixture was incubated for 2 hours at 4°C, followed by centrifugation (30 min, 11000×*g*) at 4°C. The pellet containing phage particle was resuspended in 30 ml of TSM buffer.

The MS2 RNA was purified from the phage particles using QIAamp Circulating Nucleic Acid Kit (Qiagen, Germany) accordingly to the manufacturer’s protocol and stored at −80°C.

### cDNA synthesis

The template RNA for cDNA synthesis was isolated from human HEK293 cell culture with TRIzol reagent (Thermo Fisher Scientific, USA), or FFPE samples, or synthesised *in vitro*, as described above. The synthesised RNA oligonucleotides (DNA-Synthesis, Russia) cel-miR238, cel-miR39, hsa-miR-16-5p were used as miRNA substrates (Table 1).

**Table 1.**
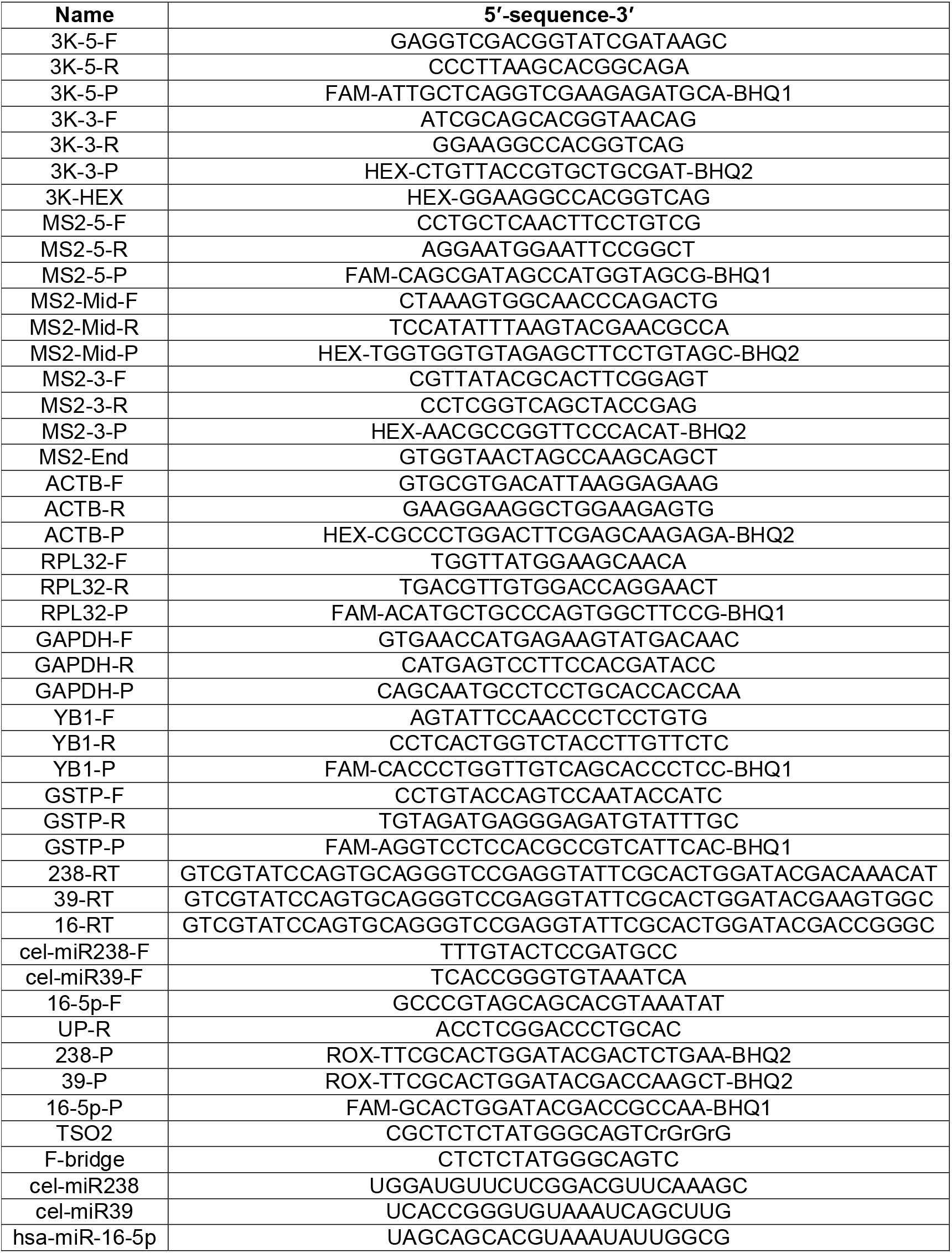
Primers and other oligonucleotides.

The 10 μL reaction probes containing 100 ng - 1μg of RNA template (or 10^6^ copies of miRNA), and 2 μM random N9, or oligo(dT)25 for RNA from HEK293 and FFPE, MS2-End for MS2 RNA, 3K-3-R for artificial RNA, 238-RT, 39-RT, 16-RT for miRNAs as primers (Table 1) were heated for 3 min at 65°C, followed by cooling on ice at 5 min. After priming, 10 μL of 2× reaction buffer (100 mM Tris-HCl (pH 8.3), 150 mM KCl, 20 mM DTT, 6 mM MgCl_2_ 0.5 mM dNTPs) with 200 U of the assayed RT were added to cooled template-primer mixtures, and the probes were immediately transferred to a preheated thermal cycler for a 20 min incubation at 37°C followed by 42°C for a 20 min, and inactivation by heating at 90°C for a 3 min. The reaction products were analysed using quantitative real-time PCR or droplet digital PCR.

Tolerance of RTs to various inhibitors was analysed by addition at the cDNA synthesis of inhibitors at various concentrations: formamide, phenol, guanidine isothiocyanate, guanidine chloride, NaCl, ethanol, urea, human whole blood, and human blood plasma (collected from five healthy individuals).

### Terminal transferase assay

The synthesised RNA oligonucleotides (DNA-Synthesis, Russia) cel-miR238, cel-miR39, hsa-miR-16-5p were used as substrates for a terminal transferase assay (Table 1).

The 6 μL reaction probes containing 10^6^ copies of synthetic miRNAs (cel-miR238, cel-miR39, hsa-miR-16-5p [Table 1]), 2 μM corresponding stem-loop primer (238-RT, 39-RT, 16-RT [Table 1]), and 2 μM oligonucleotide TSO2 (Table 1) were heated for 3 min at 65°C, followed by cooling on ice. After priming, 6 μL of 2× reaction buffer (100 mM Tris-HCl (pH 8.3), 150 mM KCl, 20 mM DTT, 6 mM MgCl_2_ 0.5 mM dNTPs) with 200 U of the assayed RT were added to cooling template-primer mixtures, and the reactions were immediately transferred to a preheated thermal cycler for a 40 min incubation at 42°C. After heating, 3 μL of 5× reaction buffer for tailing (250 mM Tris-HCl pH 8,5, 375 mM KCl, 50 mM DTT, 40 mM MnCl_2_) was added to the reactions in a thermal cycler, followed by the incubation at 42°C for 1.5 hours and inactivation by heating at 90°C for 3 min. The reaction products were analysed using quantitative realtime PCR.

### Processivity measurement

The processivity of RTs was measured using primer extension. For the RNA-dependent synthesis, the HEX-labelled primer 3K-HEX (Table 1) was hybridised to *in vitro* synthesised RNA at a 1:2 molar ratio (50 nM and 100 nM, respectively) in 1× cDNA synthesis buffer. For the DNA-template the HEX-labelled primer M13-HEX (Table 1) was hybridised to a single-stranded genomic DNA of M13mp8 phage at a 1:2 molar ratio (50 nM and 100 nM, respectively) in 1× cDNA synthesis buffer. After annealing, 5 μL of the primed template were mixed with 5 μL of the enzyme (10 U) in 1× cDNA synthesis buffer and 0.2 mM dNTPs on ice. The probes were incubated for 10 min at 37°C and quenched by the addition of 10 μL of 0.125 M EDTA. The reaction products were analysed on an ABI PRISM 3130 instrument using Peak Scanner 1.0 software (Applied Biosystems, CA). Processivity was calculated as described in Ricchetti et al. (M. and Buc H., 1993) using the following equation:

P = [[(1×I(1)]+[(2×(I(2)]+…+[(n)×(I(n))])/[I(1)+I(2)+…+I(n)]], with P being processivity, I - □the area of each peak, n□ - the number of nt added.

### Quantitative PCR analysis

The quantitative analysis of cDNA synthesis products was performed using PCR with TaqMan-probes. Each sample was analysed in triplicate. The reactions were performed in 20 μL volume containing 1×PCR buffer (65 mM Tris-HCl, pH 8.9, 24 mM (NH_4_)_2_SO_4_, 0.05% Tween-20, 3 mM MgSO_4_), 0.2 mM dNTPs, 300 nM primers, 100 nM TaqMan probe (Table 1) and 1 U of Taq-polymerase. A standard curve was generated to determine the amount of cDNA, using serial dilutions of human genomic DNA, containing multiple pseudogenes of loci, used for cDNA quantification. The amplification was carried out in CFX96 Real-Time PCR Detection System (Bio-Rad, USA) according to the following program: 95 °C for 3 min followed by 45 cycles of 95 °C for 10 s, and 60 °C for 40 s with a collection of fluorescent signals at each respective channel.

### Droplet digital PCR

The ddPCR was performed using the QX100 system (Bio-Rad, USA) according to the manufacturer’s recommendations. The 20 μL ddPCR reaction mixture contained 1× ddPCR master mix (Bio-Rad, USA), 0.9 μM primers, 0.25 μM probe (Table 1), and 40 ng of tested cDNA. The entire reaction mixture together with 70 μL of droplet generation oil (Bio-Rad, USA) was loaded into a disposable plastic cartridge (Bio-Rad, USA) and placed in the droplet generator. After processing, the droplets obtained from each sample were transferred to a 96-well PCR plate (Eppendorf, Germany). The amplification was carried out using T100TM Thermal Cycler (Bio-Rad, USA) according to the program: DNA polymerase activation at 95°C for 10 min followed by 45 cycles of PCR amplification (94°C for 30 s and 57°C for 60 s), and 98°C for 10 min, 2°C/s ramp rate at all steps. After the PCR, the droplets were counted with the QX100 Droplet Reader. The data obtained were analysed with QuantaSoft software (Bio-Rad, USA).

## RESULTS

### The composition of M-MuLV RT fusions

Here, we have obtained five chimeric enzymes based on the M-MuLV RT and DNA-binding proteins according to the cloning procedure in pET23b vector followed by an expression in *E. coli* cells of BL21 (DE3) pLysS strain. All proteins were purified using affinity and ion-exchange chromatography with 99% purity (Figure 1B). The proteins were constructed as: DBD-RT and RT-DBD – M-MuLV RT with the DNA-binding domain of *Pyrococcus abyssi* DNA ligase (DBD) fused to N- or C-terminus, respectively; Sto-RT and RT-Sto – M-MuLV RT with Sto7d fused to N- or C-terminus, respectively; RT-Sto mut – M-MuLV RT with D200N, T330P, L139P mutations and Sto7d at C-terminus. In was previously shown that Sto7d has ribonuclease activity. However, the introduction of the single-point mutation K12L disabled its RNAse activity (Shehi et al., 2001). A schematic representation of all studied proteins is shown in Figure 1A. The attachment of the DBD and Sto7d to the different parts of M-MuLV RT could provide information about the effects of additional domains on the enzymatic properties. (Moreover, Sto7d was added to a C-terminus of previously described mutant M-MuLV RT (D200N, T330P, L139P) to increase the RNA binding affinity of the enzyme.

**Figure 1.**
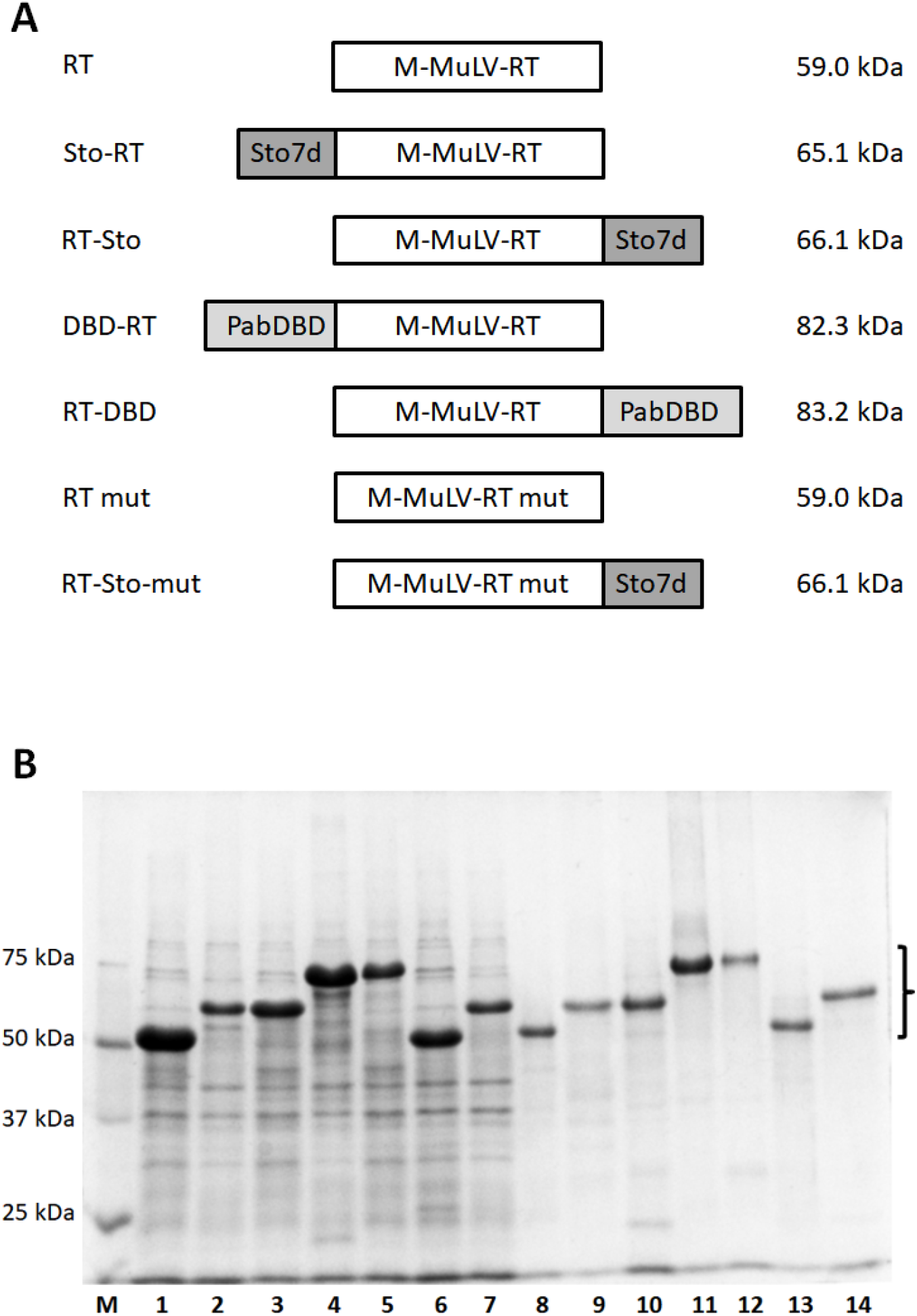
Schematic representation (A), expression, and purification (B) of M-MuLV RT fusions. RT - M-MULV RT H-, RT mut - M-MULV RT H-with D200N, T330P, L139P mutations, DBD – DNA-binding DNA-binding domain of DNA ligase *Pyrococcus abyssi*, Sto – Sto7d from thermophilic archaea *Sulfolobus tokodaii*. Enzymes were expressed in *E. coli* strain BL-21 (DE3) pLysS and purified using affinity and ion-exchange chromatography. M – Precision Plus Protein standards (Bio-Rad, USA), 1 – RT, 2 – Sto-RT, 3 – RT-Sto, 4 – DBD-RT, 5 – RT-DBD, 6 – RT mut, 7 – RT-Sto mut. Chimeric proteins are marked with a bracket.

### Additional domains do not influence the thermal denaturation of the chimeric enzymes

The thermal stability of RTs is a crucial property to realise the productivity of cDNA synthesis *in vitro*. The RNA secondary structure mostly melts at the reaction temperature range of 50-65°C, which prevents the prevents stalling of enzymatic movement and facilitates the reverse transcription in general.

The stability of the chimeric enzymes at various temperatures was investigated using differential scanning fluorimetry (DSF). The DSF utilises the ability of specific fluorescent dyes to bind with hydrophobic regions of proteins. During thermal denaturation, soluble globular proteins undergo structural rearrangement, leading to the inner hydrophobic regions being exposed out of the protein globule. The binding of the fluorescent dyes with the emerging hydrophobic surfaces leads to a detectable increase in a fluorescent signal. A shift of the thermal profile indicates the change in protein stability, and the magnitude of the change reflects the degree of folding alteration. The relevant comparative parameter is the melting temperature modulation.

The denaturation temperature of the chimeric M-MuLV reverse transcriptases did not differ significantly regardless of the type and positioning of the additional domain, or the presence of mutations (Figure 2, Table 2). The melting temperature of the original M-MULV RT was around 50°C. Only RT-Sto mut displayed lower temperature around 47°C, with all other chimeric enzymes melted at the range of 49-51°C. Notably, in the presence of primed template (poly(rA)/oligo(dT)_25_), the melting temperature of all studied enzymes remained the same (data not shown). According to the experimental results, all chimeric RTs can be divided into three groups in virtue of the basal fluorescence levels: low (RT-Sto), middle (RT-Sto mut, DBD-RT, RT, and RT mut), and high levelled (Sto-RT, and DBD-RT). However, no correlation of the basal fluorescence level with the type or localisation of the additional domain or with the presence of mutations was found.

**Figure 2.**
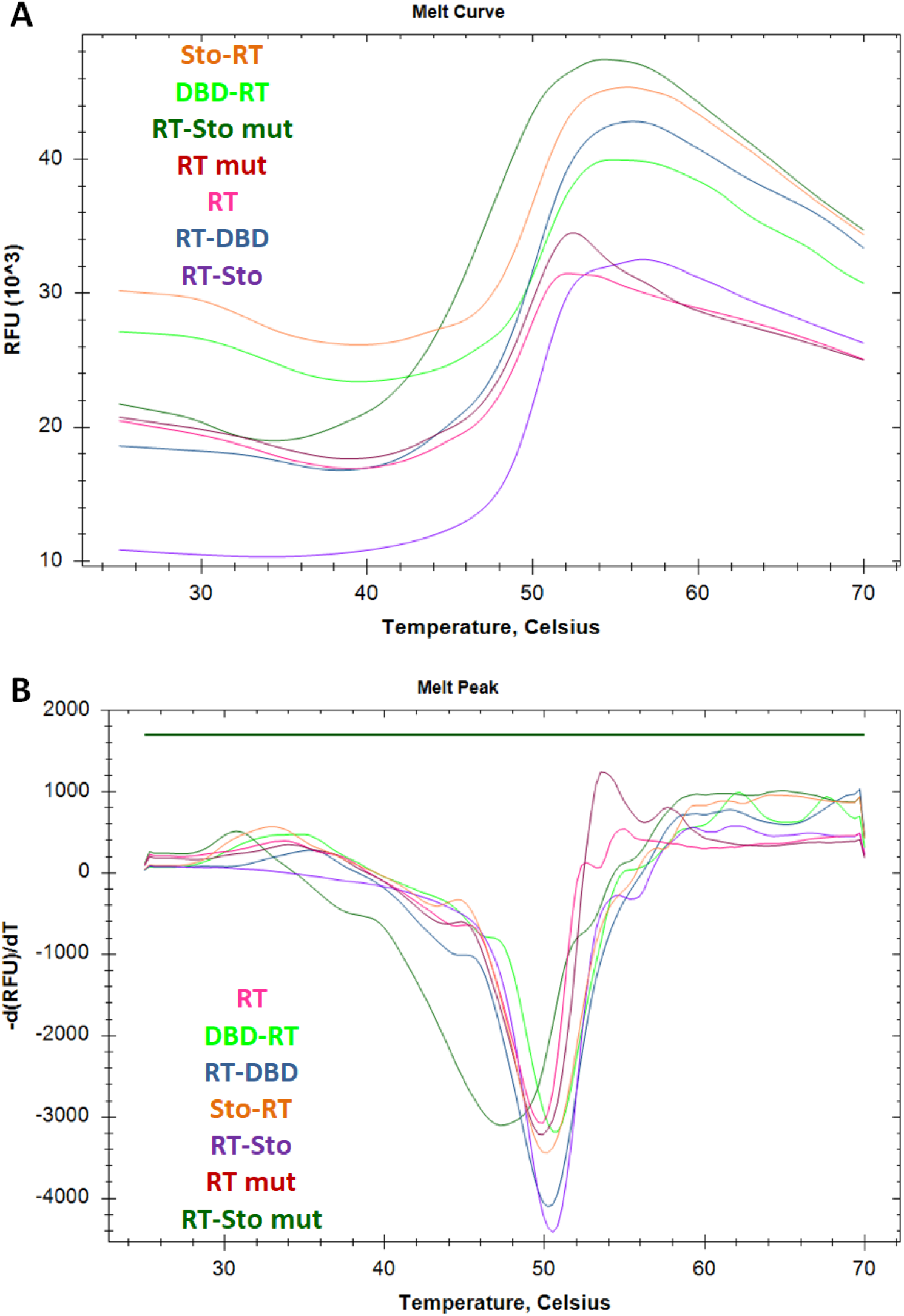
Thermal shift assay of the chimeric RTs. Thermal denaturation profiles (**A**) and the first derivative of the fluorescent curves (**B**) are presented; curve colour indicates the particular RT. Each experiment was triplicated; typical curves are presented.

**Table 2.** Specific activity, thermostability and processivity of reverse transcriptases.

| Enzyme | Specific activity, U/mg | Melting temperature, °C | Processivity, nt |  |
| --- | --- | --- | --- | --- |
|  |  |  | RNA-template<sup>1</sup> | DNA-template<sup>2</sup> |
| RT | 6,79*10<sup>4</sup> | 50.30±0.17 | 78±2 | 17±3 |
| DBD-RT | 6,99*10<sup>4</sup> | 49.38±0.45 | 82±4 | 18±5 |
| RT-DBD | 2,06*10<sup>4</sup> | 50.35±0.17 | 86±6 | 15±4 |
| Sto-RT | 2,04*10<sup>4</sup> | 49.75±0.17 | 80±7 | 18±4 |
| RT-Sto | 0.52*10<sup>4</sup> | 50.73±0.38 | 231±5 | 70±3 |
| RT mut | 9,99*10<sup>4</sup> | 50.20±0.05 | 452±6 | 80±7 |
| RT-Sto mut | 1,18*10<sup>4</sup> | 47.13±0.15 | 677±6 | 287±6 |

<sup>1</sup> - MS2 phage genomic RNA
<sup>2</sup> – M13 phage single-stranded genomic DNA

### Additional domains do not influence the temperature optimum of the chimeric enzymes

As mentioned earlier, the existence of high temperature optimum of RTs thermal stability leads to an increase in the overall efficacy of reverse transcription. To examine both characteristics of the enzymatic reaction of the chimeric proteins, we used polymerase activity assay based on the incorporation of a radioactively labelled nucleotide into poly(rA)/oligo(dT)_25_ substrate.

Firstly, we evaluated the molar activity of the RTs (Table 2). The presence of DBD at the C-end or Sto7d at N- or C-terminus of the wild-type and mutated forms of RT led to a decrease in the specific enzyme activity. The triple mutant RT mut demonstrated increased polymerase activity.

The temperature optimum for wt-based RTs was independent of the type and position of the additional domain (Figure 3A). However, as reported earlier, the mutations D200N/T330P/L139P increased the parameter up to 45°C for both RT mut and RT-Sto mut.

**Figure 3.**
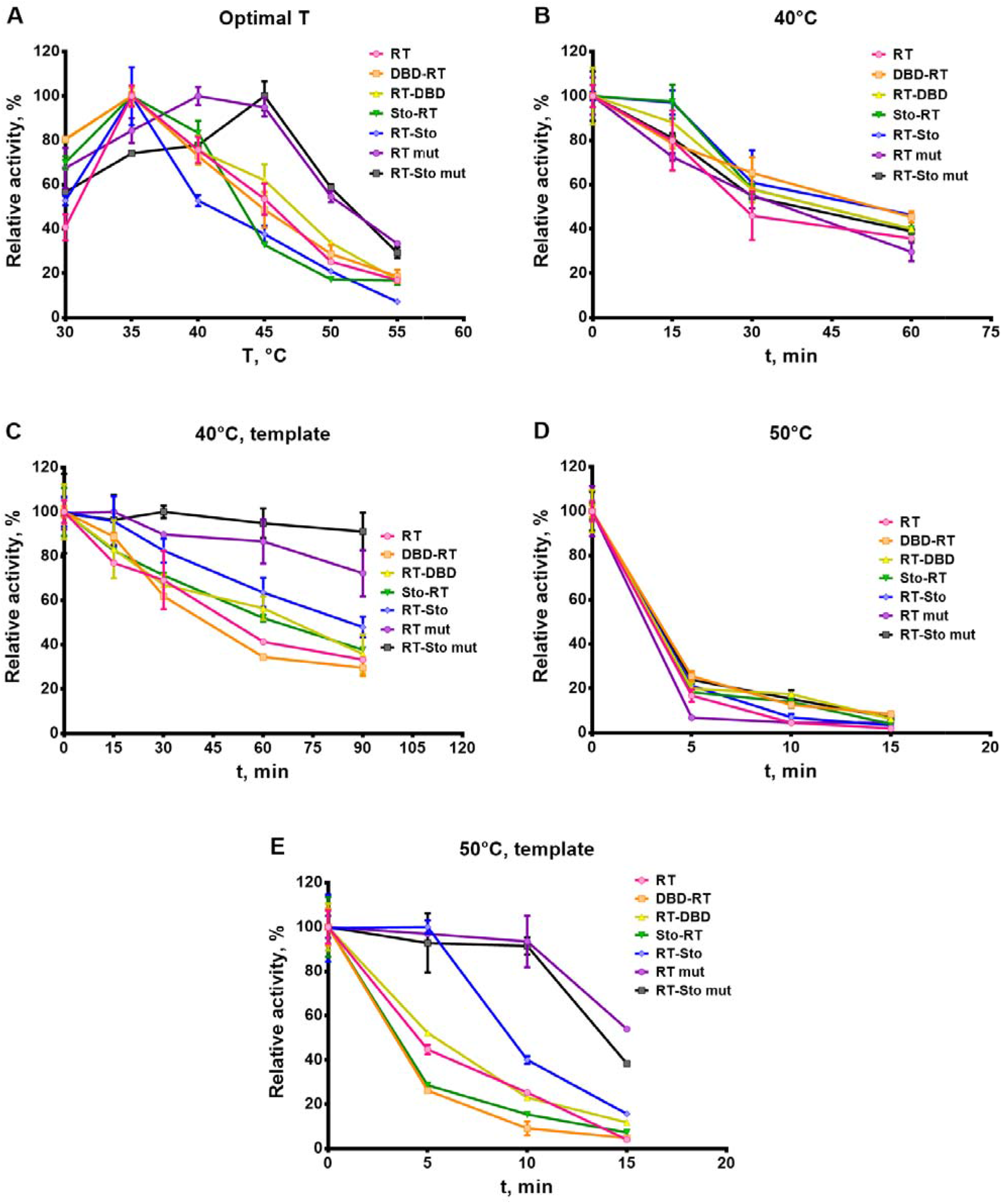
Optimal temperature (A), and thermostability of the chimeric RTs at 40 (B, C) and 50°C (D, E) in the presence (C, E) or absence (B, D) of the RNA substrate (poly(rA)/oligo(dT)_25_). Enzymatic characteristics were determined via incorporation of α-[P]-dTMP in poly(rA)/oligo(dT)_25_ substrate. Each experiment was triplicated. Optimal temperature and thermal stability were determined in standard polymerase activity assay after the incubation at indicated conditions.

The most intriguing results were obtained when studying the enzymatic ability to synthesise cDNA after prolonged heating in the presence of poly(rA)/oligo(dT)_25_, i.e. thermal stability. All the enzymes studied showed similar thermal stability, maintaining around 40% of activity after heating for 60 minutes at 40°C (Figure 3B) and around 5% – after 5 minutes at 50°C (Figure 3D). If the RNA template was added in the heating reaction, the thermal stability of all enzymes was increased, contradicting the results of the thermal stability assay (Figure 3C, 3E). The chimeric enzymes bearing mutations (RT mut and RT-Sto mut) were the most stable, with the presence of Sto7d enhancing the effect. Thus, RT mut retained around 80% of activity after 1.5 hours at 40°C, RT-Sto mut - 100%. After being heated for 15 minutes at 50°C, the RT mut retained about 40% of the initial activity, with the RT-Sto mut maintaining 50%.

### Addition of Sto7d at C-terminus increases processivity of the enzyme

Processivity is an enzyme property that, in the case of reverse transcriptases, is defined as a number of nucleotides incorporated during the synthesis into the nascent DNA strand per single binding act of an enzyme with an RNA template. High processivity facilitates further cDNA analysis due to a decrease in the amount of error number at sequencing.

The processivity of the chimeric enzymes was evaluated using the elongation of a fluorescent-labelled primer annealed to RNA or DNA templates. Despite the localisation, DBD did not affect the processivity of the respective chimeric RTs in comparison to unmodified RT (Table 2). At the same time, Sto7d increased the processivity of RT and RT mut at trifold by being attached to the C-end of these enzymes. M-MuLV RT with D200N/T330P/L139P mutations showed an increase in processivity by about 6 times compared to the original enzyme. The attachment of Sto7d to the C-end of RT mut resulted in further up to 8-fold growth of the parameter.

The processivity of the chimeric enzymes was also assessed by their ability to produce long cDNA fragments. Thus, *in vitro* RNA molecule based on Lambda phage genomic DNA sequence was used as a template for cDNA synthesis. The cDNA obtained was quantified using droplet digital PCR with two sets of primers and TaqMan probes for both 5’- and 3’-ends of the cDNA (Figure 4). The ratio of 5’-to 3’-end concentrations serves as a surrogate marker of RT processivity: more processive enzymes are able to synthesise longer cDNA molecules, resulting in a 5’/3’-ends ratio close to 100%. The results obtained correspond to the primer extension analysis: the D200N/T330P/L139P mutations and Sto7d at C-terminus increased the ability of the chimeric enzyme to synthesise long (3 kb) cDNA stretches. The original M-MuLV RT as well as DBD-RT, RT-DBD, and Sto-RT were able to elongate only 1-2% of all cDNA molecules up to the 5’-end of RNA template. The RT-Sto, RT mut, and RT-Sto mut fully synthesised around 40, 60, and 80% of all cDNA molecules, respectively.

**Figure 4.**
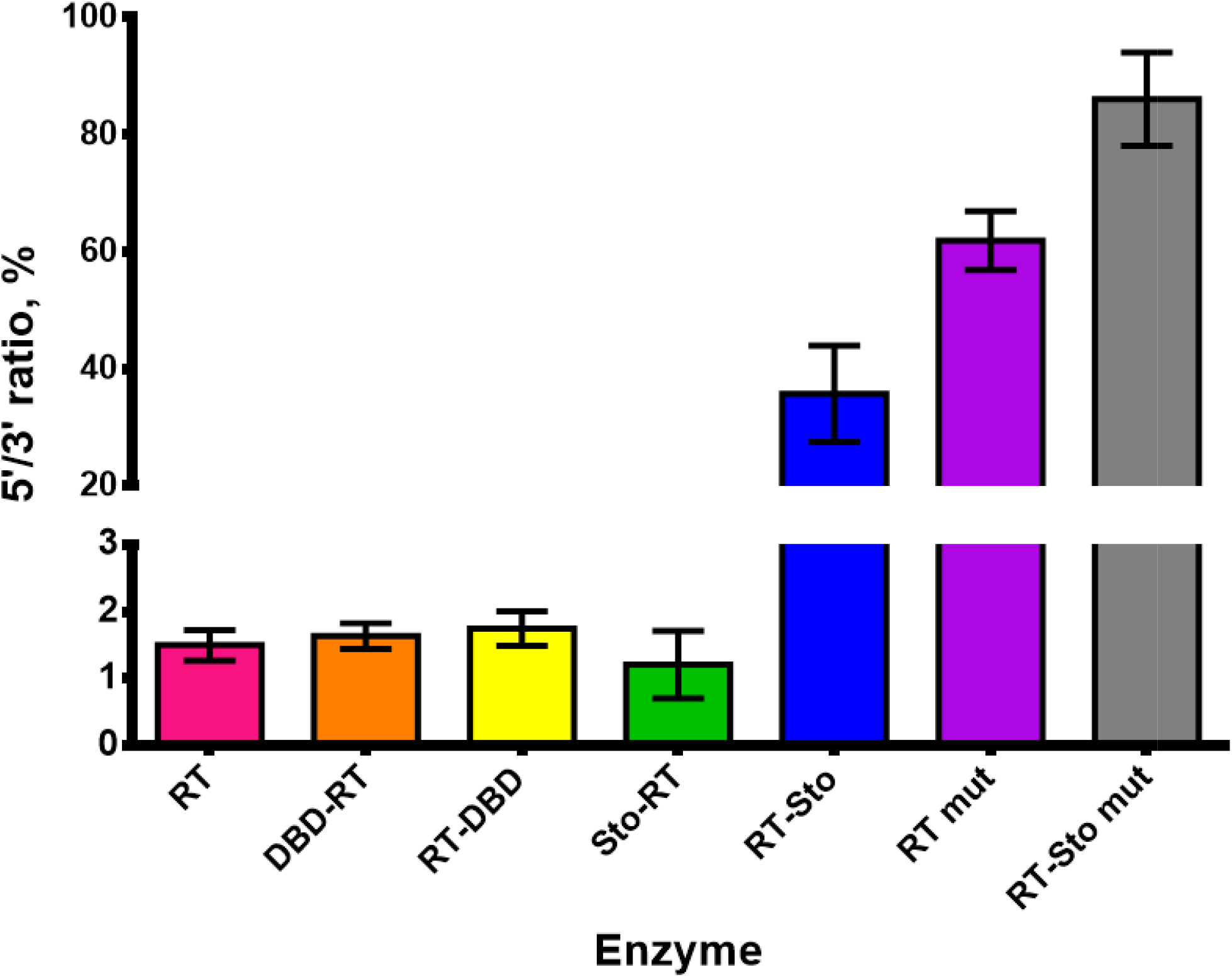
The ratio of the 5-’ to 3’-end concentrations of cDNA synthesized by RTs using the 3 kb synthetic RNA template. Each reaction was run in three technical repeats. The resulting ratio of cDNA 5- and 3’-ends was determined using ddPCR.

Reverse transcriptases have both RNA-dependent and DNA-dependent DNA-polymerase activities. The ability to produce DNA using DNA as a template is applied for the synthesis of the cDNA second strand. Thus, when preparing NGS-libraries, the RT and RNAse H would be used for double-stranded cDNA synthesis without adding DNA polymerases.

The processivity of the chimeric RTs was studied by the elongation of a fluorescent-labelled primer annealed with DNA template (T able 2). While DNA-directed processivity of enzymes with DBD and N-terminal Sto7d remained at the level of the original M-MuLV RT, C-terminal Sto7d and mutations increased the parameter considered in the same manner as when the RNA-dependent processivity was studied. The most processive enzyme was the RT-Sto mut (an 8-fold increase compared to M-MuLV RT), fully corresponding to the results of RNA-directed DNA-synthesis.

### Sto7d broadens tolerance of the enzymes to ions

The ionic strength of the solution strongly affects the electrostatic interactions of proteins with nucleic acids, which play a significant role in the binding of proteins with DNA or RNA. High salt concentrations indicate high ionic strength, with ions acting as a “screen” to prevent the interaction of charged biomolecules. Additional nucleic acids-binding domains allow respective proteins to tether DNA/RNA despite a high concentration of ions.

We tested whether the chimeric enzymes retain polymerase activity in the presence of high concentrations of mono- or divalent ions (K^+^, Na^+^, Cl^-^, SO_4_^2-^), which are present in most of the biological substances (Figure 5). The assay was based on the incorporation of a radioactively labelled nucleotide into poly(rA)/oligo(dT)_25_ substrate. It turned out that RT-Sto, RT mut, and RT-Sto mut were more resistant to all kinds of the investigated ions than the original M-MuLV RT or their counterpart with DBD. The effect of D200N/T330P/L139P mutations was exceeded by Sto7d fusion and in the RT-Sto mut having a wider salt concentration resistance range than the RT mut.

**Figure 5.**
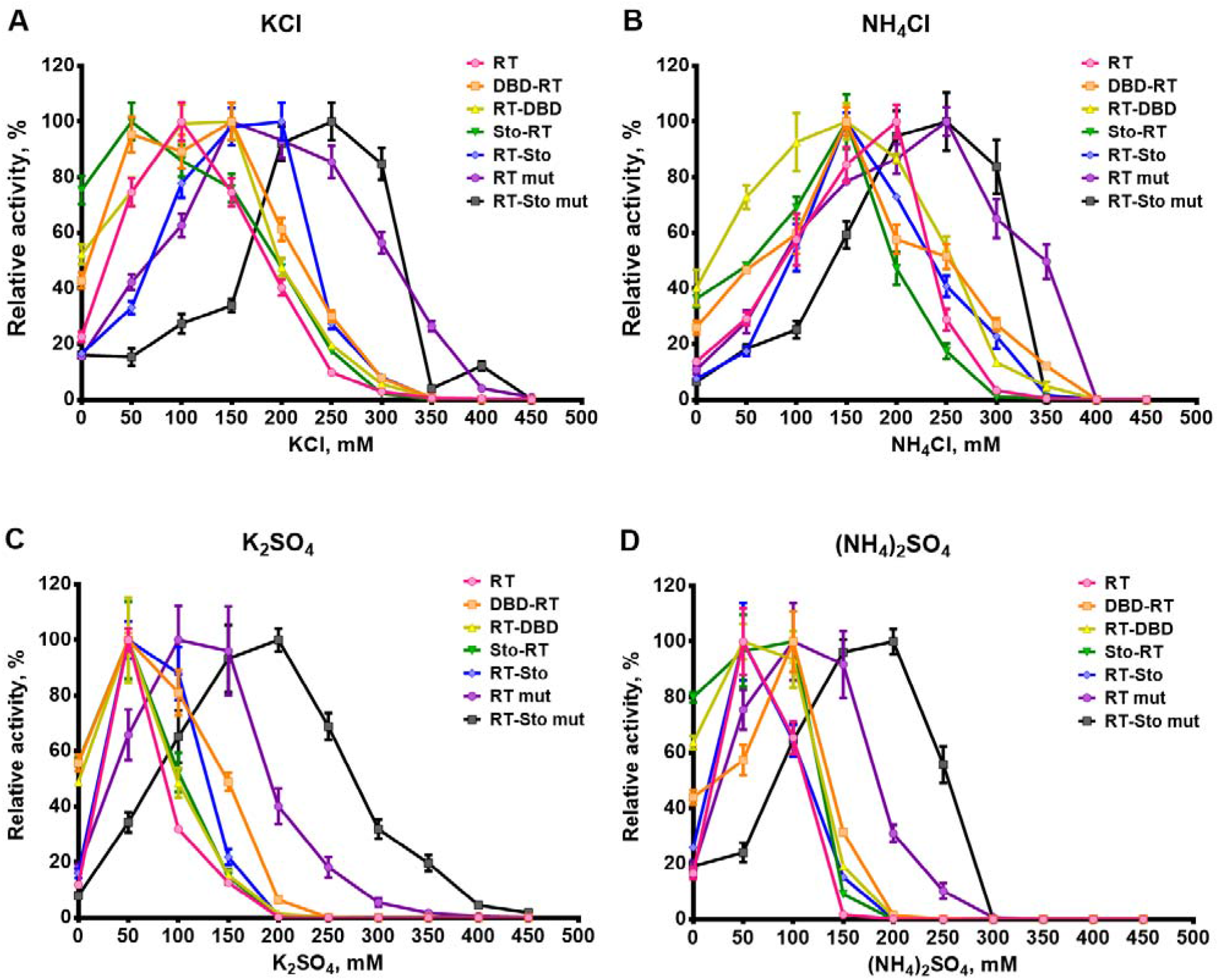
Optimal ion concentrations for the chimeric RTs. Enzyme characteristics were determined via incorporation of α-[^32^P] dTMP in poly(rA)/oligo(dT)_25_ template. Each experiment was triplicated.

### Additional domains do not affect terminal transferase activity

Terminal transferase activity, or tailing, is an ability of RT to incorporate several nucleotides into the nascent DNA strand in a template-independent manner. Tailing is used in several practical applications, including single-step preparation of cDNA fragments for NGS-analysis.

The terminal transferase activity of the RTs was studied by the assay with template-switching oligonucleotides (Figure 6A). At the first step, the hybridisation with the stem-loop-containing RNA oligonucleotide as a primer was used for cDNA synthesis. When RT synthesised cDNA up to 5’-end of the RNA template, the enzyme started to add up to 3 unpaired nucleotides to a 3’-end of the cDNA fragment; predominantly, dCMP was incorporated among other dNMP. Template-switching oligonucleotide (TS-oligonucleotide) with 3’-(r)GMP_n_-end hybridised to the 3’-CMP-overhang and serves as a template for a DNA-directed DNA synthesis. At the end of a tailing, the RT synthesised the DNA fragment consisting of the stem-loop primer, cDNA, dCMP-overhang, and DNA complementary to TS-oligonucleotide. The products of tailing reaction were quantified using qPCR with two sets of primers and probe: one universal reverse primer hybridised to a former stem-loop primer, probe hybridised to a cDNA, and two different forward PCR-primers. One forward primer (inner) was complementary to the 3’-end of the cDNA, while another forward primer (outer) – to the 3’-end of the DNA complementary to TS-oligonucleotide. The ratio of DNA concentrations obtained by using outer to inner primer indicates a tailing efficacy. The results of the assay are presented in Figure 6B.

**Figure 6.**
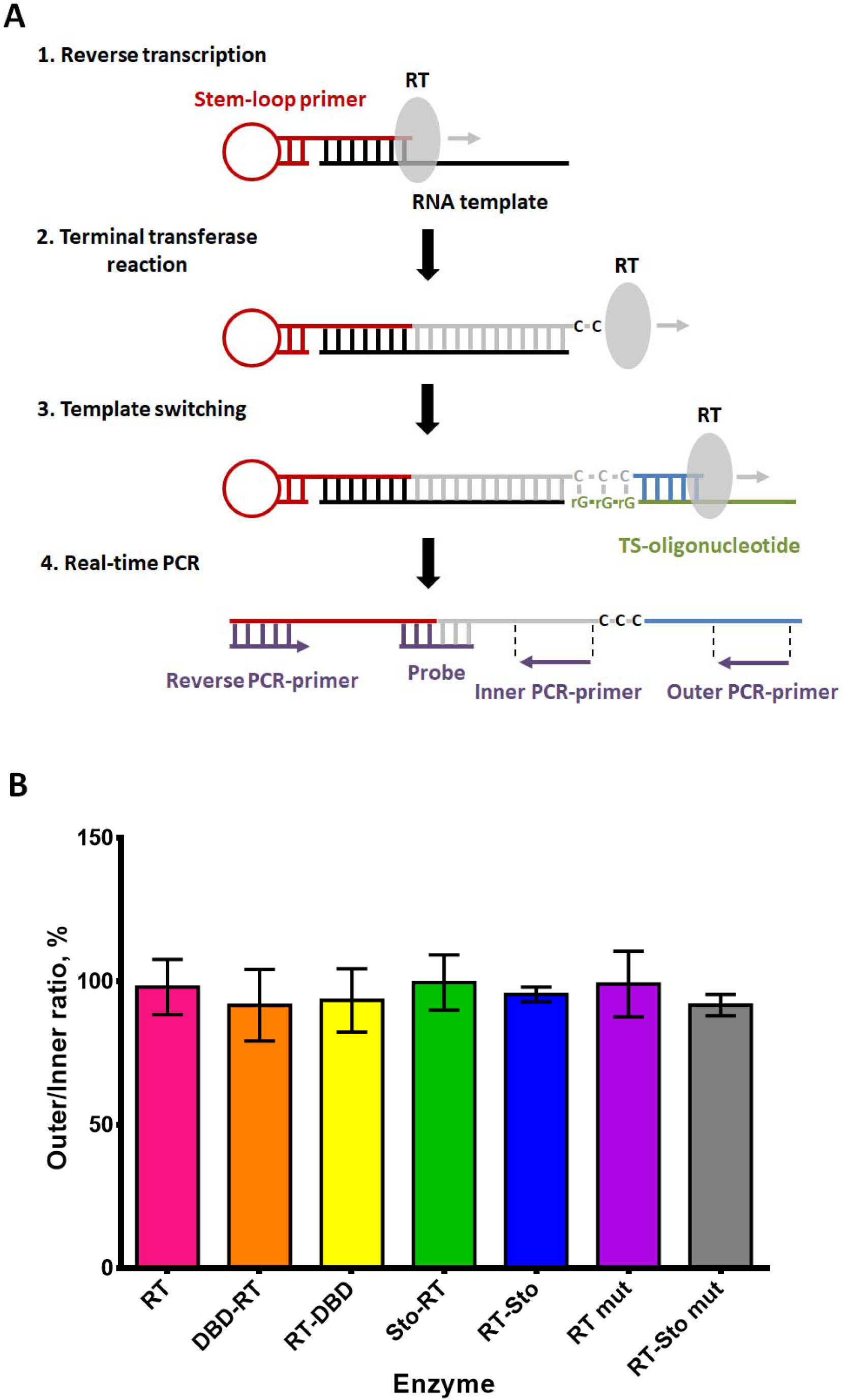
Terminal transferase activity of the chimeric RT. Scheme of the experimental procedures (**A**) and the respective results (**B**). Each reaction was carried out in three technical repeats. The resulting ratio of elongated DNA fragments after terminal transferase reaction to converted RNA template was determined using qPCR.

All chimeric enzymes retained terminal transferase activity and were able to elongate cDNA with almost 100% efficacy. Thus, all RTs are suitable for preparing the NGS cDNA libraries with TS-oligonucleotides.

### Sto7d increases the efficacy of chimeric enzymes in cDNA synthesis

The high yield of cDNA simplifies RNA-of-interest detection or quantitative and qualitative analysis of all RNA molecules in the specimen. Therefore, the amount of synthesised cDNA defines the feasibility of the specific RT for practical usage. An ability of the chimeric RTs to synthesise cDNA was studied using RNA from two sources: HEK293 cells culture, and FFPE colon cancer specimens. Additionally, three artificial miRNA served as templates for reverse transcription. Thus, we compared the efficacy of RTs on either intact (cell culture) or degraded chemically modified (FFPE) RNA-templates. With random primers providing an indiscriminating synthesis of large pools of cDNA, priming with N9 was used, while oligo- (dT)_25_ as primers allow converting only polyadenylated RNA molecules. The resulting cDNA amount was quantified with qPCR with 5 target loci (location, amplicon length and GC-content listed in Table 3).

**Table 3.** Loci for WGA quantification.

| **Gene** | **Localization** | **Length, bp** | **GC-content** |
| --- | --- | --- | --- |
| *ACTB* | 7p22.1 | 174 | 59.77 |
| *GAPDH* | 12p13.31 | 215 | 57.21 |
| *GSTP1* | 11q13.2 | 143 | 54.44 |
| *RPL32* | 3p25.2 | 74 | 48.65 |
| *YBX1* | 1p34.2 | 83 | 55.42 |

Using random N9 primers resulted in higher production of cDNA; the effect was observed for the *RPL32, ACTB*, and *YB1* genes by both types of RNA template (Figure 7). The RT mut and RT-Sto mut outperformed other enzymes for the most loci with all combinations of template and primers; The RT-Sto mut was superior to the RT mut. RT-Sto synthesised less cDNA amount than the RTs with the mutations. Among the extant enzymes, DBD-RT outperformed M-MuLV-RT, RT-DBD, and Sto-RT that were similar. The efficacy of miRNA conversion to cDNA by the studied RTs had an identical level (Figure 8).

**Figure 7.**
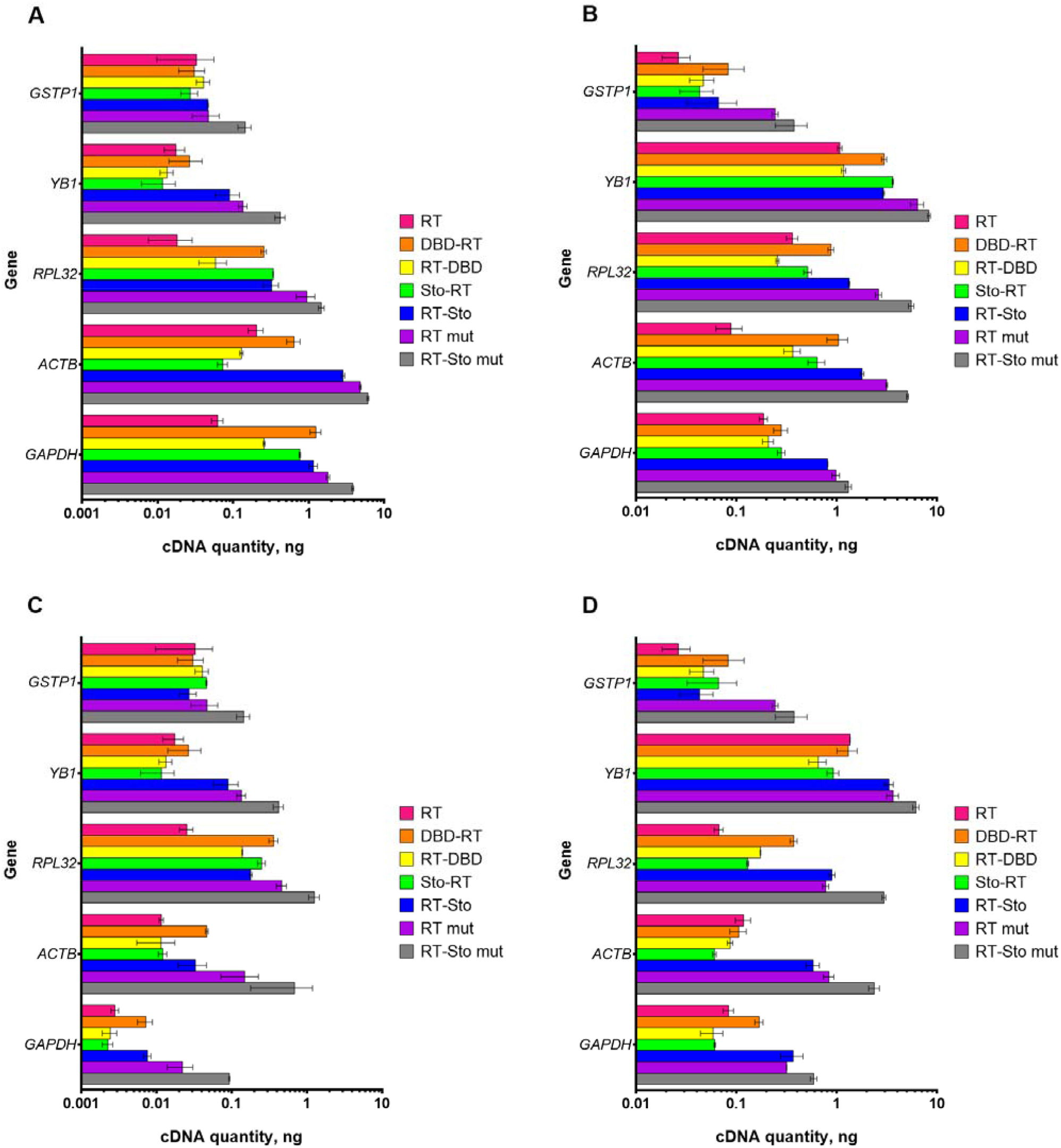
Synthesis of cDNA by the chimeric RTs using a total RNA as a template. Each synthesis reaction was run in triplicate. Total RNA from HEK293 cells (A, B), or total RNA from FFPE samples (C, D) were used as the template, primed by oligo-(dT)_25_ (A, C), or N9 (B, D). A standard curve was generated to determine the amount of produced cDNA for each locus.

**Figure 8.**
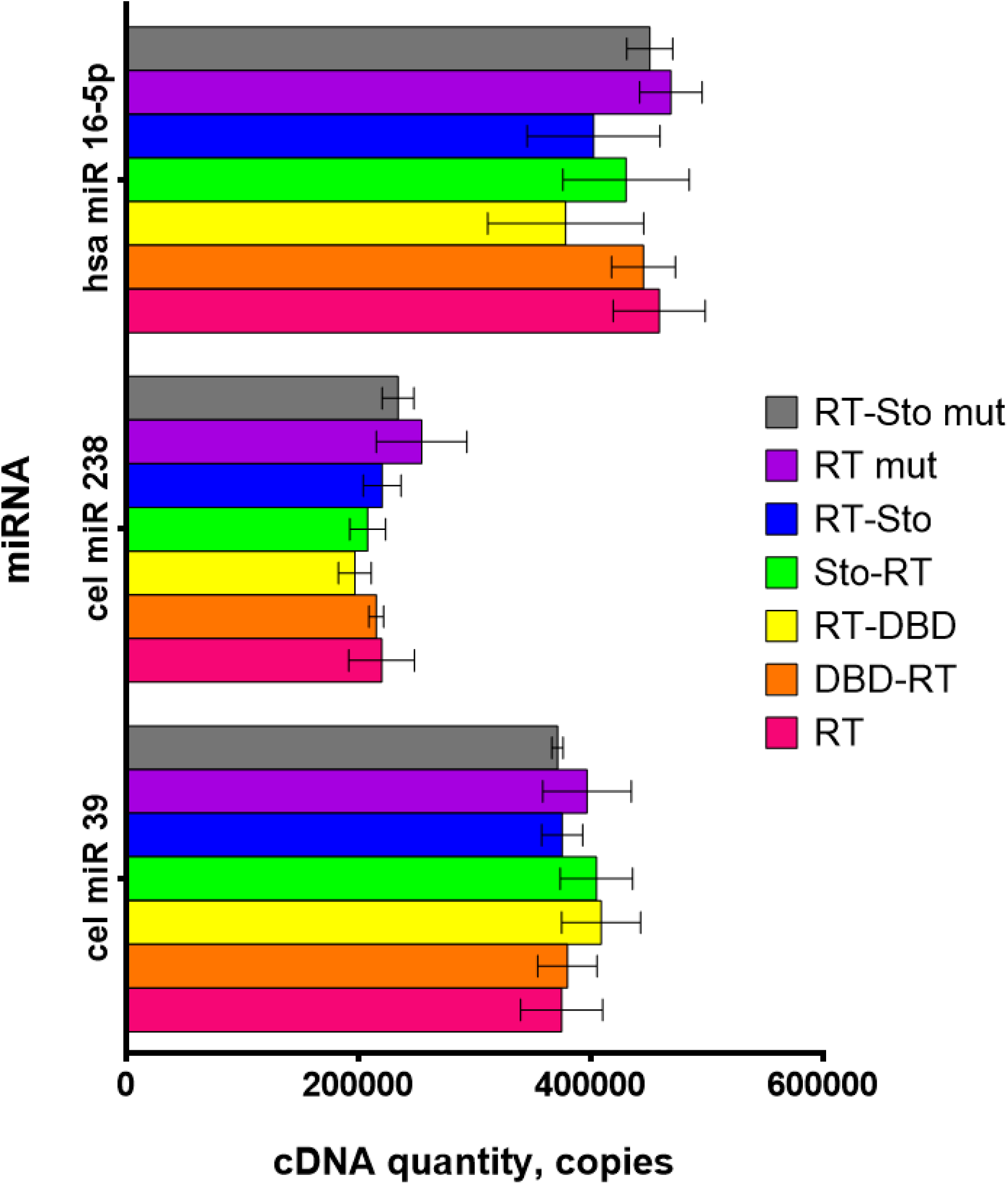
Synthesis of cDNA with the chimeric RTs using the short RNA templates. Each synthesis reaction was run in triplicate. Artificially synthesised RNA oligonucleotides, mimicking miRNAs, served as the templates. A standard curve was generated to determine the amount of produced cDNA for each locus.

### Sto7d increases tolerance to inhibitors

RT is an indispensable component of multiple test-systems intended for the detection of human and agricultural pathogens such as retroviruses, viroids. In terms of diagnostics, the RT’s tolerance to inhibitors is one of the significant factors contributing to overall test robustness. This parameter is of great importance for developing the point-of-care approach in diagnostics when simple tests are needed, and DNA/RNA purification is impracticable.

Here, we tested the tolerance of the chimeric enzymes to common inhibitors arising from biological substances or reagents from purification procedures: ethanol, urea, phenol, formamide, NaCl, guanidinium chloride, and guanidinium isothiocyanate. The inhibitors were added in different concentrations into reaction mixtures for the reverse transcription, followed by a qPCR analysis of the resulting cDNA products (Figure 9).

**Figure 9.**
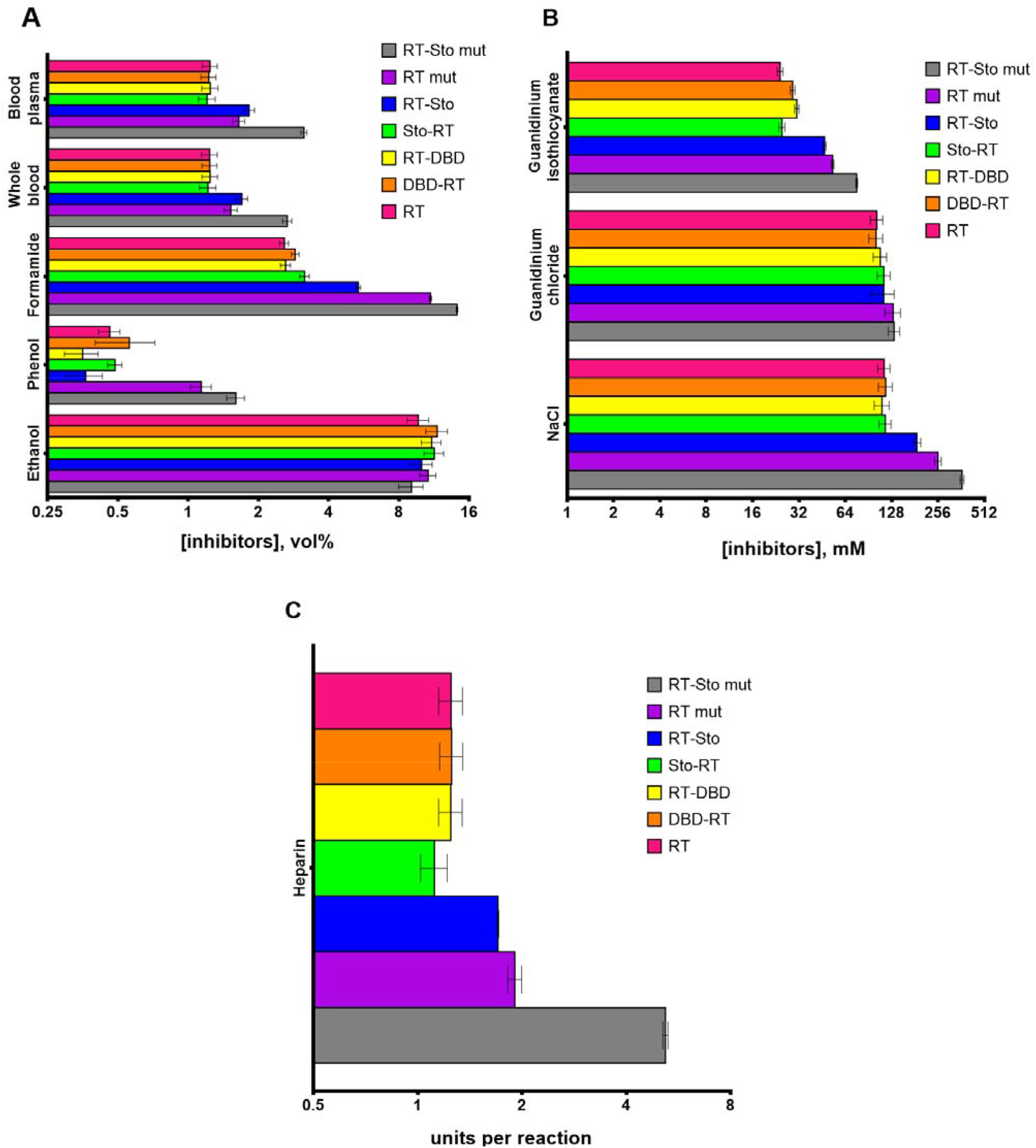
Tolerance to inhibitors of the chimeric RTs. Substances tested as inhibitors in a cDNA synthesis: ethanol, phenol, formamide, human whole blood, human blood plasma (A), NaCl, guanidinium isothiocyanate, guanidinium chloride (B), heparin (C). Each cDNA synthesis reaction was run in triplicate. The artificial 3 kb transcript was used as the template. IC50 values for different inhibitors were calculated and plotted on the graphs.

The tolerance to NaCl of RT-Sto, RT mut, and RT-Sto-mut was increased, while the tolerance to ethanol and guanidinium chloride remained the same for all the enzymes. The presence of either triple mutations or Sto7d enhanced the ability of RTs to synthesise cDNA in the presence of formamide and guanidinium isothiocyanate. In both cases, the effect of mutations and Sto7d was summed. The same distribution was observed under inhibition by urea: RT-Sto, RT mut, and RT-Sto-mut were insusceptible to 2 M urea, while other enzymes worked up to 1.5 M urea (data not shown). The more complicated situation was obtained under investigation of phenol effect: RT mut and RT-Sto mut were more robust, while RT-Sto activity remained at the level of the original RT. However, RT-Sto mut surpassed RT mut in the means of tolerance to phenol. In the case of complex biological substances (e.g., human whole blood and human blood plasma), the obtained results were contradicted to previously described. C-terminal Sto7d amended the effect of the inhibitors more efficiently than triple mutations. However, RT-Sto mut was the most productive.

## DISCUSSION

For the past few decades, RTs have become an indispensable tool for academic researchers and laboratory diagnosticians. Multiple studies have been made to obtain more thermostable and processive enzymes. Most researchers have focused on discovering new enzymes or introducing single-point mutations into RTs described, especially into commonly used M-MuLV and ALV RTs. Previously, fusion with additional domains was efficiently applied for constructing DNA polymerases with improved processivity and tolerance to various inhibitors. Thus, small RNA- and DNA-binding proteins from Sso7d-like family were attached to DNA polymerases from family A (Taq-polymerase (Wang et al., 2004), Sau-polymerase (Zhai et al., 2019), Gss-polymerase (Oscorbin et al., 2017)), family B (Pfu-polymerase (Wang et al., 2004), Tgo-polymerase (Jozwiakowski and Connolly, 2011), KOD-polymerase (Wang et al., 2015)), and family Y (Dbn-polymerase (Wu et al., 2017)). Helix-hairpin-helix (HhH) domains from thermostable enzymes were also attached to several DNA polymerases (de Vega et al., 2010; Pavlov et al., 2002). However, to our knowledge, no similar fusion RTs have been made based on M-MuLV RT, or AVL RT. In the present study, we intended to fill the gap and construct novel enzymes with improved properties.

A set of chimeric RTs was constructed based on M-MuLV RT and two DNA-binding domains: DNA-binding domain of DNA ligase *Pyrococcus abyssi* and DNA-binding protein Sto7d from thermophilic archaea *Sulfolobus tokodaii*. Additional DNA-binding domains were fused with either N- or C-end of M-MuLV RT, making it possible to take into account possible adverse effects of the added domain in the specific localisation. Previously described as increasing the processivity, optimal temperature, and tolerance to inhibitors the mutations D200N, T330P, L139P were added to M-MuLV RT to compare their effect with the effect of additional domains.

When studied by DSF, the melting temperature of RTs, reflecting the thermal stability of the proteins, remained the same irrespectively of the presence of mutations or additional domains. This observation is in contradiction with the previously noted fact of a change in thermostability upon a fusion of Gss-polymerase with the same DNA-binding domains (Oscorbin et al., 2017), as well as with increased optimal temperature of the mutated RTs (RT mut, RT-Sto mut). The presence of the primed RNA template also did not result in an increase in the melting temperature. It is inconsistent with the increased thermostability of RTs in the presence of the primed RNA substrate, observed in the present and past works (Gerard et al., 2002). Several explanations for the observed contradiction can be considered. The first is a possible inadequate sensitivity of DSF, which may be insufficient for detecting the minuscule conformation transitions of RTs during heat denaturation. The second is the absence of a direct connection between the increased ability of RTs to bind the template and structural changes during heating, which also could be a result of the absence of a direct interplay between the RT and an additional domain. The third is maintaining the active site conformation while the whole protein globule undergoes massive structural rearrangements. In that case, primed RNA substrate may play an important role in maintaining an active site conformation, protecting it from disruption during heating. In either case, conflicting results of DSF and functional analysis of polymerase activity emphasise the necessity to design the experiments and interpret the data obtained carefully.

Additional domains did not influence the optimal temperature of the RTs, while the mutations increased the parameter up to 45°C, as it was reported (Arezi and Hogrefe, 2009). At the same time, adding Sto7d at C-end of both original M-MuLV RT and RT mut increased the heating time before the loss of polymerase activity. Notably, the effect was observed only in the presence of a primed RNA template, implying the stabilisation of the enzymes by binding with the RNA. RT-Sto, RT mut, and RT-Sto mut also demonstrated the increased tolerance to a salt concentration and processivity when using either RNA- or DNA-template. It should be noted that the characteristics of the fusions with DBD and Sto-RT remained at the level of the original M-MuLV-RT. While the ability of the DBD to bind with RNA is unknown, similar HhHs are widespread among the proteins involved in both DNA and RNA metabolism (Shao and Grishin, 2000). Gss-polymerase with DBD at the N-end showed increased processivity when using a DNA template (Oscorbin et al., 2017). However, the processivity of DBD-RT on the DNA template was similar to M-MuLV RT. The latter implicates on the unpredictability of interactions between noncognate protein domains, thereby making it unreliable to directly transfer results from the one fusion protein to another one with a similar architecture. The increased processivity of RT-Sto, RT mut, and RT-Sto mut can facilitate the synthesis of long cDNA molecules. A small length of cDNA fragments hampers the analysis of NGS data, making the assembly of sequence reads a cumbersome task, and increasing the possibility of “bad mapping”. Longer cDNA molecules allow avoiding the problem; therefore, processive RTs (RT-Sto, RT mut and RT-Sto) are more suitable for NGS-libraries preparation.

The main focus of previous studies was to increase the thermostability and the optimal temperature of different RT. While numerous successful attempts resulted in stable and processive RTs (Arezi and Hogrefe, 2009; Baba et al., 2017; Baranauskas et al., 2012b; Ellefson et al., 2016; Heller et al., 2019; Michael J. Moser et al., 2012; Sauter and Marx, 2006; Zhao et al., 2018b), other vital RT characteristics were not clarified in several studies (Arezi et al., 2010b). Thus, high tolerance to inhibitors is crucial for the development of robust tests for point-of-care (POC) diagnostics. POC diagnostics is intended for the testing at the patient’s bedside when DNA purification procedures should be simplified or omitted, resulting in different cellular and chemical contaminations in the probe. Some of them, e.g. heparin, blood and blood plasma components (hem, lactoferrin, bile salts, bilirubin, IgG) come from the biological substances being analysed. The others (EDTA, phenol, guanidinium salts, NaCl, urea, ethanol) are the components of buffers meant for nucleic acids purification. In either case, the presence of inhibitors may lead to false-negative results of testing and subsequent inadequate therapy (Acharya et al., 2017; Hedman and Rådström, 2013; Whiley et al., 2006). The attachment of Sto7d not only improved the processivity of M-MuLV RT but also increased the tolerance to chaotropic agents, disturbing protein-nucleic acids interactions. RT-Sto mut was also able to synthesise cDNA at an increased concentration of human whole blood and human blood plasma which was in accordance with the similar effect observed for chimeric Gss-polymerase with Sto7d. The fact supports the hypothesis of compromising the protein-RNA or protein-DNA interaction by unknown components of human blood.

Due to its DNA-dependent DNA polymerase and strand-displacement activity, RT is able to synthesise the cDNA second strand independently of other enzymes. A terminal transferase activity, the ability to incorporate nucleotides into nascent cDNA strand in a template-independent manner, provides the opportunity for producing double-stranded cDNA without attracting an external DNA polymerase. While the specific activity of the chimeric RTs was affected by the additional domains, the terminal transferase activity remained intact. Thus, the chimeric enzymes could be used for a DNA-polymerase-free preparation of NGS cDNA libraries.

The ability of RT to synthesise cDNA using RNA template with a complex secondary structure or high GC-content is one of the major characteristics defining the usability for practical applications. In viral RNA, the specific secondary structures can participate in the regulation of replication, transcription and translation. At 45-50°C RNA secondary structure could hamper cDNA synthesis, causing a systematic bias in transcriptome assay data or false-negative results of the virus diagnostics. The type of priming also influences the cDNA product quantity and quality. Using random primers may lead to a spurious amplification and unnecessary amplification of ribosomal RNAs while using oligo(dT) primers may produce a lesser amount of cDNA due to an RT stalling at the secondary structures of non-unfolded RNA templates. The attachment of Sto7d at C-terminus of M-MuLV-RT resulted in a higher amount of cDNA, independent of the priming method. The ability to synthesise cDNA using complicated RNA template, such as MS2 phage genomic RNA, was also increased by the chimeric enzymes with Sto7d.

Several concerns should be noted as a subject for future studies. For example, proteins with higher affinity to RNA could be more advantageous for constructing the chimeric RTs. RNA-binding domains of nucleocapsid proteins from several viruses or SSB protein from the hyperthermophilic archaeon *Sulfolobus solfataricus* showed tether binding to RNA (Chang et al., 2014; Morten et al., 2017). However, excessively strong coupling with the template may hinder catalysis or lead to cell toxicity, hampering the induction of the enzyme. The fidelity of the chimeric RTs described here remains unstudied. However, in some cases, including testing of somatic mutations, the fidelity of RT becomes a crucial parameter to be investigated thoroughly. In theory, an additional domain should not affect the structure of the active site. Moreover, natural high processive polymerases tend to be more accurate. Last but not least, other enzymes with reverse transcriptase activity will also be considered as an appealing target for fusing with additional domains.

To sum up, we have proved that it is possible to improve RT efficiency by attaching an additional domain. The chimeric M-MuLV RT with Sto7d showed an enhanced ability to synthesise cDNA, as well as increased robustness to common RT inhibitors.

## ACKNOWLEDGEMENT

The authors express profound gratitude to Ekaterina A. Belousova for the invaluable help in the preparation of the manuscript.

