## Supplementary file for "The attachment of Sso7d-like protein improves processivity and resistance to inhibitors of M-MuLV reverse transcriptase"

**Construction of M-MuLV reverse transcriptase fusions**

***RT***

The M-MuLV reverse transcriptase nucleotide sequence was amplified using RT-F1/RT-R1 primers (Supplementary Table) with NheI and NotI restriction sites, allowing the in-frame ligation into the pET23b vector (Novagen, USA). PCR was carried out using as template M-MuLV cDNA. The resultant 1.7-kbp DNA fragment and pET23b vector were digested with NdeI and NotI (SibEnzyme, Russia), ligated, and transformed into *E. coli* XL1-Blue cells according to the standard protocols (26). The fidelity of the resulting recombinant plasmid named pET-RT was confirmed by sequence analysis using primers pET-F and pET-R (Supplementary Table).

***RT mut***

The plasmid pUC-RT contained partial coding sequence M-MuLV reverse transcriptase with mutations D200N, T330P, L139P was constructed by Shanghai RealGene Bio-tech, Inc (China). Plasmid pUC-RT was digested with KpnI/SalI, and DNA fragment coding mutated M-MuLV RT was eluted from agarose gel, ligated with pET-RT (KpnI/SalI), and transformed into *E. coli* XL1-Blue cells according to the standard protocols. The resulting plasmid was named pET-RT-mut.

***DBD-RT fusion***

Pab-DBD and partial M-MuLV-RT nucleotide sequences were amplified using DBD-F1/DBD-R1 and RT–F2/RT-R2 primers (Supplementary Table), respectively. PCR was carried out using previously constructed pET-DBD (27), containing the nucleotide sequence of the DBD of ATP-dependent DNA ligase from *Pyrococcus abyssi*, and pET-RT. Resultant PCR fragments were digested with BamHI, followed by ligation according to the standard protocols. The fusion DNA fragment was eluted from agarose gel, digested with NdeI/KpnI, and ligated with pET-RT (NdeI/KpnI). The resultant plasmid was named pET-DBD-RT.

***RT-DBD fusion***

The Pab-DBD and partial M-MuLV-RT nucleotide sequences were amplified using DBD-F2/DBD-R2 and RT-F3/RT-R3 primers (Supplementary Table), respectively. PCR was carried out using previously constructed pET-DBD, and pET-Gss. Resultant PCR fragments were fused via PCR with RT-F3/DBD-R2 primers. The fusion DNA fragment was eluted from agarose gel, digested with SalI/NotI, and ligated with pET-RT (SalI/NotI). The resultant plasmid was named pET-RT-DBD.

***Sto-RT fusion***

The Sto7d and partial M-MuLV-RT nucleotide sequences were amplified using Sto-F1/Sto-R1 and RT-F2/RT-R2 primers (Supplementary Table), and the resulting DNA fragments were fused via PCR with Sto-F1/RT-R2 primers. PCR was carried out using previously constructed pET-Sto-Gss (24), containing the mutated nucleotide sequence of the Sto7d from *Sulfolobus tokodaii,* and pET-RT. Resultant PCR fragments were fused via PCR with Sto-F1/RT-R2 primers. The fusion DNA fragment was eluted from agarose gel, digested with NheI/KpnI, and ligated with pET-RT vector (NheI/SalI). The resultant plasmid was named pET-Sto-RT.

***RT-Sto fusion***

The Sto7d and partial M-MuLV-RT nucleotide sequences were amplified using Sto-F2/Sto-R2 and RT-F3/RT-R4 primers (Supplementary Table). PCR was carried out using previously constructed pET-Sto-Gss, and pET-RT. Resultant PCR fragments were fused via PCR with RT-F3/Sto-R2 primers. The fusion DNA fragment was eluted from agarose gel, digested with SalI/NotI, and ligated with pET-RT vector (SalI/NotI). The resultant plasmid was named pET-RT-Sto.

***RT-Sto mut fusion***

The Sto7d and partial M-MuLV-RT nucleotide sequences were amplified using Sto-F2/Sto-R2 and RT-F3/RT-R4 primers (Supplementary Table). PCR was carried out using previously constructed pET-Sto-Gss, and pET-RT mut. Resultant PCR fragments were fused via PCR with RT-F3/Sto-R2 primers. The fusion DNA fragment was eluted from agarose gel, digested with SalI/NotI, and ligated with pET-RT vector (SalI/NotI). The resultant plasmid was named pET-RT-Sto-mut.

Supplementary table. Primers for cloning of the chimeric RTs.

| **Name** | **5′-sequence-3′** | **Restriction site** |
|---|---|---|
| RT-F1 | TATG**GCTAGC**CTAAATATAGAAGATGAGCATCGGC | NheI |
| RT-R1 | GAGT**GCGGCCGC**ATCAAGGCAGTTGTGTTGC | NotI |
| DBD-F1 | TCATG**CATATG**AGGTACATAGAGCTGGCCCA | NdeI |
| DBD-R1 | ATTC**GGATCC**CTTTATTGGCTTACCAATCTGAATT | BamHI |
| RT-F2 | ATTC**GGATCC**ctaaatatagaagatgagcatcggc | BamHI |
| RT-R2 | GATGATGGTACCAGTATTCCCTGGTCC | KpnI |
| DBD-F2 | ATTCAGATTGGTAAGCCAATAAAGAGGTACATAGAGCTGGCCCA | |
| DBD-R2 | GATGAT**GCGGCCGC**ATTAGCTAATCCATCATTACCCTCA | NotI |
| RT-F3 | CTCTTT**GTCGAC**GAGAAGCA | SalI |
| RT-R3 | CTTTATTGGCTTACCAATCTGAATATCAAGGCAGTTGTGTTGC | |
| Sto-F1 | GTCTC**GCTAGC**ATGGTAACAGTAAAGTTCAAGTATAA | NheI |
| Sto-R1 | GCCGATGCTCATCTTCTATATTTAGACCGCCACCGCCTTTCTTTCCAGATTTTTCTAACATTT | |
| Sto-F2 | GGTACCGGCGGTGGCGGTGTAACAGTAAAGTTCAAGTATAA | |
| Sto-R2 | GATGAT**GCGGCCGC**TTTTTCTAACATTTGTAGTAATTCTT | NotI |
| RT-R4 | ACCGCCACCGCCGGTACCATCAAGGCAGTTGTGTTGC | |
| pET-F | CCTATAGTGAGTCGTATTAATTTC | |
| pET-R | CAACTCAGCTTCCTTTCGG | |
